# Shared Molecular Landscape of Human Brain Structure and Systemic Metabolism

**DOI:** 10.1101/2025.01.12.632645

**Authors:** Dennis van der Meer, Jakub Kopal, Sara E. Stinson, Olav B. Smeland, Alexey A. Shadrin, Julian Fuhrer, Jaroslav Rokicki, Tiril P. Gurholt, Ida E. Sønderby, Ibrahim A. Akkouh, Oleksandr Frei, Kevin S. O’Connell, Lars T. Westlye, Srdjan Djurovic, Anders M. Dale, Ole A. Andreassen

## Abstract

The human brain is metabolically demanding, yet the causal interplay between systemic metabolism and brain morphology remains largely unknown. Applying systematic phenome- and genome-wide analyses of 24,940 UK Biobank participants, integrating 249 NMR-derived plasma metabolic markers with MRI-derived brain morphology, we revealed extensive phenotypic associations, with 236 of 249 metabolites linked to at least one global brain measure, including intracranial volume (192 markers), total surface area (183), and mean cortical thickness (139). Distinct regional brain patterns included frontal, temporal, brainstem, and ventricular structures, and the inflammatory marker glycoprotein acetyls showed the strongest individual association, explaining the most variance in global brain measures. We identified extensive genetic overlap, encompassing genes involved in neurodevelopment and stem cell differentiation. Mendelian randomization revealed significant bidirectional causal effects, with metabolic markers predominantly influencing frontal and temporal brain regions. Single cell sequencing data implicated oligodendrocytes and astrocytes mediating the relationships. These findings highlight extensive links between metabolic processes and brain structure, and provide an atlas for understanding the underlying molecular pathways.

---

The human brain’s high energy requirements demand a tight regulation of nutrient supply to enable its development, functioning, and maintenance.^1^ This intensive metabolic activity underpins the brain’s continuous integration of peripheral metabolic signals to modulate both central and systemic metabolic activity and maintain homeostasis.^2^ Accumulating evidence highlights the critical role of systemic metabolism in brain development and disease.^3^ Brain disorders are associated with distinct plasma metabolite profiles^4–6^ and an increased burden of cardiometabolic disease, which severely impacts morbidity and mortality.^7,8^ However, the precise causal relationships between systemic metabolic activity and brain health remain incompletely understood; complexity arises from multifaceted interactions, including metabolic exchange across the blood-brain barrier (BBB),^9^ neurohumoral signaling, and immune-mediated communication pathways. Given the growing global burden of brain and cardiometabolic disorders in the aging population,^10^ it is imperative to resolve this knowledge gap to better understand these pressing public health challenges.

Metabolomics and neuroimaging data are complementary modalities that can capture lifespan pathogenic processes across the brain-body axis. High-throughput nuclear magnetic resonance (NMR) spectroscopy allows for comprehensive quantification of plasma metabolic markers, including lipoproteins, cholesterol, fatty acids, amino acids, and glucose,^4^ beyond traditional measurements, often limited to a handful of lipid markers.^11^ These metabolite classes are highly relevant to brain development and function. Indeed, metabolic programming during critical neurodevelopmental windows profoundly influences brain structure and lifelong functional trajectories.^12^ The brain consumes 25% of the body’s energy derived from glucose, primarily to maintain proper ion gradients and neurotransmitter levels at synapses.^13^ While both neurons and astrocytes are glucose-dependent, astrocytes can additionally supply neurons with lactate via metabolic coupling,^14^ highlighting a dynamic intercellular metabolic network. Lipids, comprising 50% of brain dry weight,^15^ are essential for development and maintenance of brain structure and function,^16^ facilitating cell differentiation, signal conductance, and synaptic throughput.^17,18^ Further, amino acids are fundamental for neurotransmitter biosynthesis and modulation,^19^ directly impacting synaptic function. Metabolic dysregulation, driven by factors such as insulin resistance and systemic inflammation, can cause mitochondrial dysfunction, oxidative stress, and neuroinflammation within the brain,^20^ ultimately impacting cellular viability and brain morphology.

Small sample sizes and limited biomarker coverage constrained prior investigations of metabolite-brain morphology relationships. Nevertheless, evidence links specific markers, including high- and low-density lipoproteins (HDL, LDL),^21–23^ glutamate, glutamine,^24^ and inflammatory markers,^25^ to regional brain structures. We and others have previously linked a range of cardiometabolic risk factors to macro-scale brain structure, with recent work extending beyond body mass index (BMI)^26^ to specific body composition and inflammatory markers.^27^ Post-mortem studies of individuals with psychiatric disorders have further shown abnormal metabolic marker levels in the prefrontal cortex, striatum and thalamus.^28–30^

Genomics offers a powerful approach to clarify causal relationships and shared biology between systemic metabolism and brain morphology. Mendelian randomization (MR) studies have indicated causal effects of individual circulating lipids on cortical morphology.^31,32^ Extensive genetic overlap also exists between cardiometabolic diseases and brain disorders.^33,34^ Using genome-wide association studies (GWAS), we and others have mapped the genetic architecture of metabolic markers^35,36^ and brain morphology,^37,38^ identifying thousands of genetic associations.

Despite accumulating evidence, a large-scale systematic investigation of shared molecular pathways between *in vivo* human brain morphology and systemic metabolism has been lacking. To address this gap, we leveraged data from 24,940 UK Biobank (UKB) participants, integrating 249 NMR-derived plasma metabolite concentrations with 91 MRI-derived measures of cerebral cortical thickness (TH), surface area (SA), and subcortical volumes. We additionally integrated large-scale GWAS data from over 200,000 individuals to systematically assess causal pathways and shared genetic architecture. We report strong, widespread phenotypic and genetic associations between plasma metabolic markers and brain morphology. Our analyses identify regional patterns of association, provide evidence for bidirectional causal relationships, and reveal shared molecular pathways implicated in neurodevelopment and stem cell differentiation. This work offers novel insights into the fundamental relationship between systemic metabolism and brain structure, and the resulting atlas offers a rich resource for identifying mechanistic targets for interventions to preserve brain health across the lifespan.^39^

## Results

Our study sample included 24,940 UK Biobank participants with both NMR metabolomics and T1 brain MRI data (mean age at blood collection=55.2 (SD=7.9); mean age at brain scan=64.0 years (SD=7.5) 52.5% female). We quantified brain morphology using Freesurfer to derive 88 regional measures (34 regional TH and SA measures,^40^ 20 subcortical volumes), and 3 global measures (intracranial volume (ICV), total SA, mean TH).^41^ The NMR data comprised 249 plasma metabolic markers from the Nightingale panel,^4^ encompassing 228 lipids, lipoproteins or fatty acids and 21 non-lipid traits (e.g., amino acids, inflammation-related markers),^35^ see Supplementary Tables 1 and 2. For each analysis, we used the Benjamini-Hochberg correction for multiple comparisons (□=.05) across the markers and brain measures. Models were pre-residualized for age at baseline and age at scan, sex, scanner, and scan quality and rank-based inverse normal transformed. Regional brain measures were additionally corrected for their metric-specific global measure.

### Associations between brain morphology and metabolic markers

We began by testing the overarching hypothesis that metabolism, as an integrated system captured by circulating markers, is associated with normal variation in brain morphology. For this, we first applied partial least squares (PLS) regression. This revealed significant associations between the full set of metabolic markers and each global brain measure, most strongly ICV and SA (**Fig. 1a**). Subsequent linear regressions revealed an extensive link with global brain measures, with 236 of the 249 metabolic markers (94.8%) showing a significant association with at least one global brain metric. This included numerous individual associations for ICV (192 markers), SA (183 markers), and TH (139 markers). Among regional measures, temporal lobe thickness, along with brainstem, ventricular, and ventral diencephalon volumes, showed the highest number of significant associations (**Fig. 1b**).

**Figure 1:**
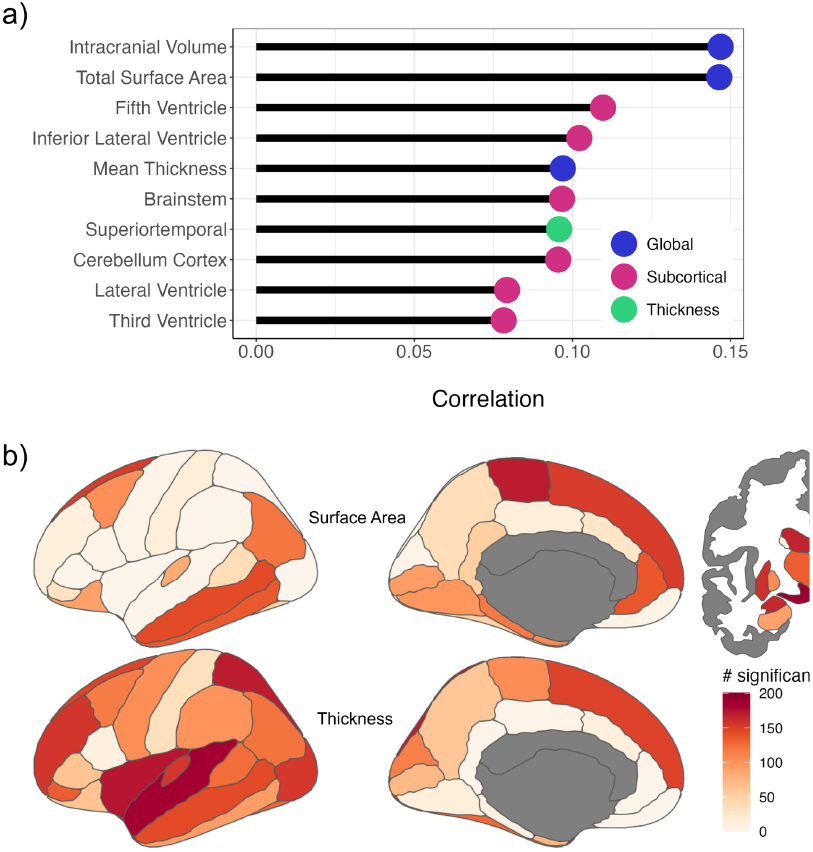
Widespread phenotypic associations link metabolic markers to brain morphology. **a)** Top 10 brain measures (y-axis) showing the strongest joint correlation (x-axis), as the square root of explained variance, with the full set of 249 metabolic markers, identified by partial least squares (PLS) regression. Colors denote the brain metric category. **b)** Regional brain maps showing the total number of significant associations (color-coded) between the 249 metabolites and cortical thickness, surface area, and subcortical volumes.

At the individual marker level, associations were strongest with ICV and total SA, whose regression coefficients across the 249 markers were highly correlated (r=0.94), while their correlations with mean TH were lower (**Fig. 2a**). The inflammatory marker glycoprotein acetyls (GlycA) had the strongest single association, explaining 1% of variance in both ICV and SA. Many markers showed strong, spatially varying associations, as exemplified by GlycA (**Fig. 2b**; full maps in Supplementary Fig. 1 and statistics in Supplementary Table 3). These findings were robust to the exclusion of individuals with a neurodegenerative disease diagnosis (n=269), with regression coefficients remaining highly correlated (r=0.95) and near-identical median absolutes (|β|=0.0107 vs 0.0106; Supplementary Table 4).

**Figure 2:**
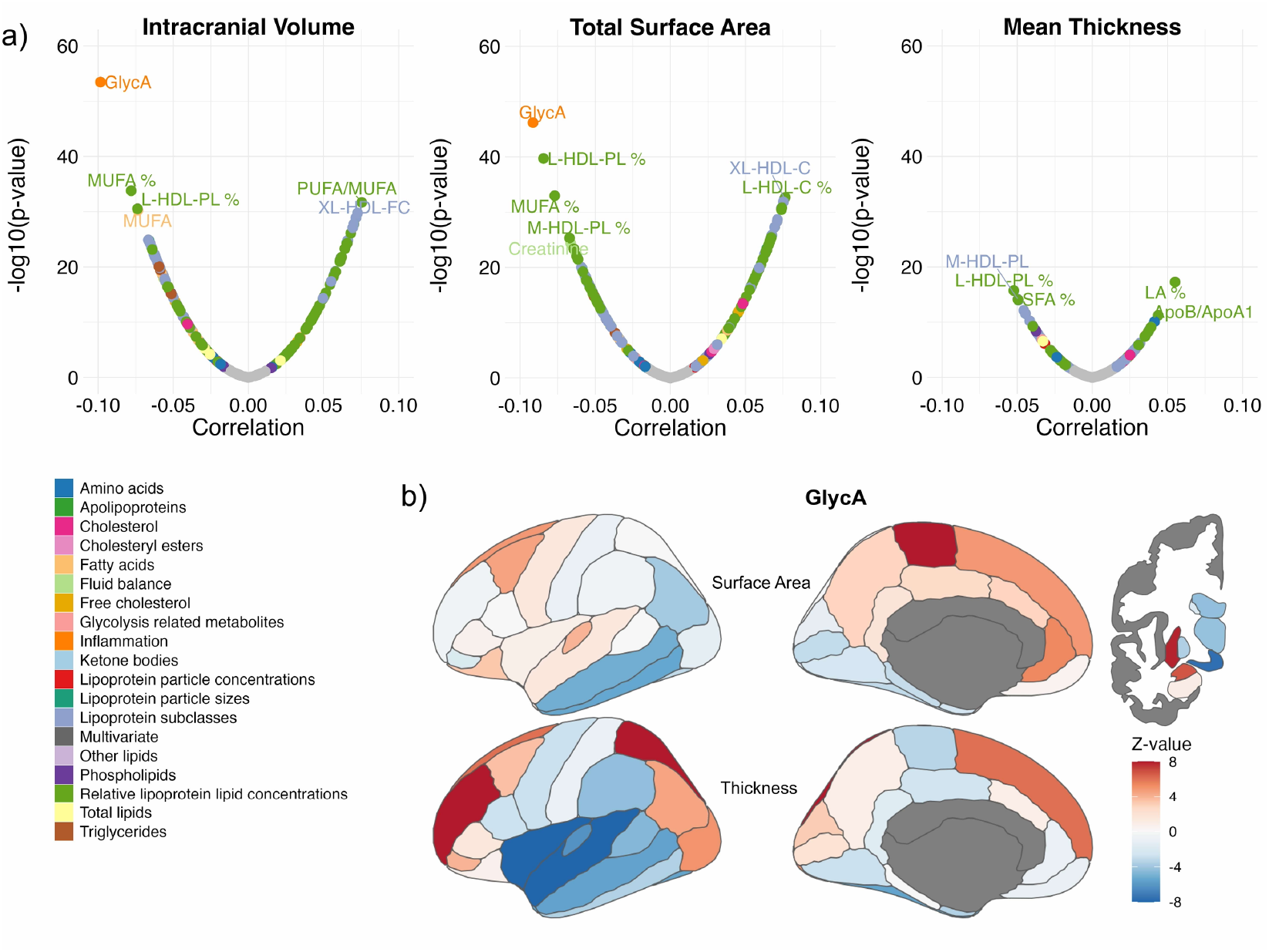
Individual metabolic markers show widespread and spatially specific associations with brain morphology. **a)** Association summary plot for each of the 249 metabolic markers with global brain measures. The x-axis shows the correlation and the y-axis shows the significance as -log10(P-value). Dots are colored by metabolite category, and top associations are annotated by name. **b)** Brain maps showing the spatial pattern of association (z-value) for one exemplar marker, glycoprotein acetyls (GlycA), with regional brain measures.

### Canonical correlations between brain morphology and metabolic markers

Next, to determine whether the observed associations reflected coherent multivariate patterns rather than isolated links, we applied canonical correlation analysis (CCA). We identified two primary components of shared variance between metabolites and brain morphology. The first component (r=0.33) primarily captured global brain measures, with high positive loadings for SA and ICV. For markers, it was driven by opposing loadings from HDL (positive) and GlycA (negative) (**Fig. 3**, left column). The second component (r=.21) revealed a pattern of larger ventricles and smaller subcortical volumes with lower LDL concentrations (**Fig. 3**, right column). Full feature loadings are provided in Supplementary Table 5.

**Figure 3:**
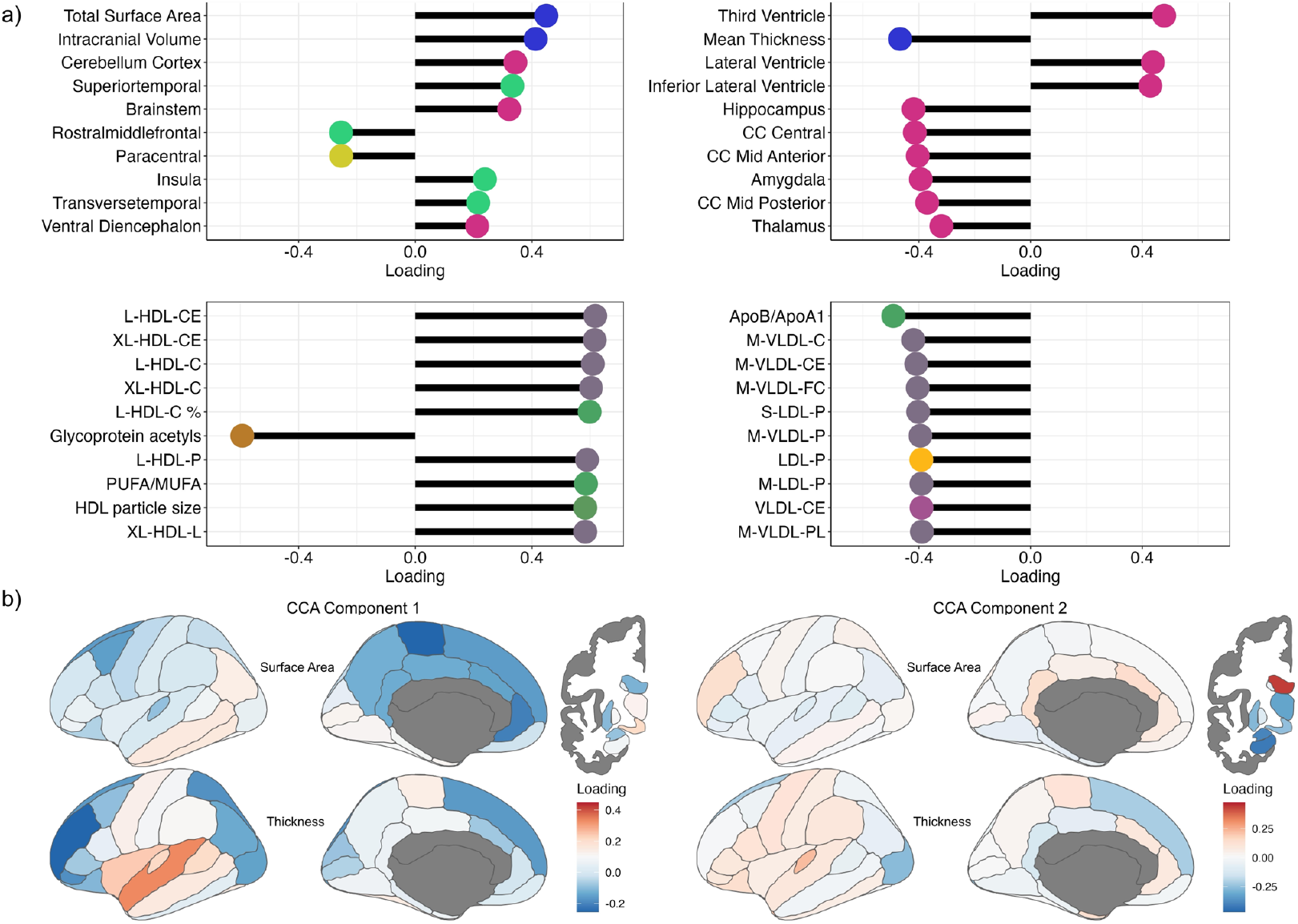
Canonical correlation analysis reveals shared structure between metabolites and brain morphology. Loadings for the top two canonical components are shown. **a)** Bar plots of the ten brain measures (top) and metabolic markers (bottom) with the highest loadings (x-axis) for each component. Marker colors distinguish brain metric types and metabolite categories. **b)** Brain maps showing the spatial distribution of loadings for each component across regional brain measures. The left column displays results for component 1; the right column displays results for component 2.

### Causal relationships between metabolic markers and brain morphology

We then examined potential causal effects between metabolism and brain morphology using bidirectional MR. This revealed widespread causal effects of metabolites on brain morphology (Supplementary Table 6). The most significant effects involved the cerebellum, with omega-6 fatty acids and alanine showing strong negative causal impacts on cerebellar cortex volume (β=-0.12, p=4.8×10^−13^; β=-0.17, p=8.13×10^−11^). We also identified robust lipid-brain interactions, including a negative influence of medium-density LDL cholesterol on thalamic volume (β=-0.28, p=9.6×10^−9^) and widespread effects of various lipoproteins and fatty acids on surface area and thickness of specific frontal and temporal cortical regions (**Fig. 4a)**.

**Figure 4:**
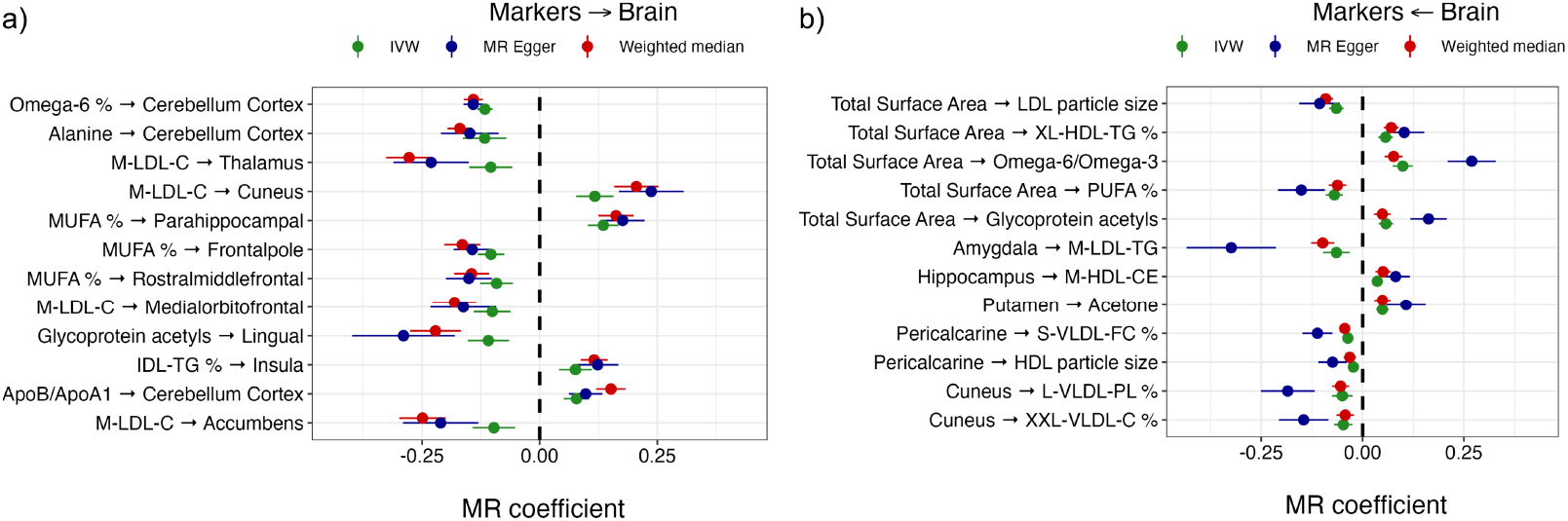
Bidirectional causal relationships between metabolic markers and brain morphology. Forest plots of illustrative causal estimates from Mendelian randomization (MR) analyses, with **a)** listing effects of metabolites on brain morphology, and **b)** causal effects of brain morphology on metabolites. The x-axis shows the causal estimate (β) with standard error, and dot color indicates the MR method used.

Furthermore, our analyses identified significant causal influences in the opposite direction, from brain morphology, primarily SA, to systemic metabolite levels (**Fig. 4b**). Notably, larger SA was causally associated with lower levels of saturated fatty acids (β=-0.08, p=2.1×10^−5^). Cross-referencing both analyses revealed 20 bidirectional relationships. For instance, we observed an antagonistic loop between small LDL cholesterol and cuneus thickness, where higher LDL was causal for a thicker cuneus (p=.001), which in turn was causal for lower LDL levels (p=.0004), suggesting a potential complex regulatory feedback loop.

### Global genetic overlap

To determine whether shared genetic architecture underlies the identified relationships between metabolomics and brain morphology, we quantified genetic architectures using Gaussian mixture modeling (MiXeR). Metabolic markers and brain measures showed comparable SNP-based heritability (median vs.13.17), while the polygenicity of metabolic markers was substantially smaller than that of brain measures (median 326 vs. 1,764 causal variants; Supplementary Fig. 2, Supplementary Table 7). Next, we estimated genome-wide genetic correlations using LD score regression (LDSC). The patterns of genetic correlation largely mirrored the phenotypic associations, with strong relationships for ICV and SA with GlycA and PUFA/MUFA (the ratio of polyunsaturated to monounsaturated fatty acid; **Fig. 5a**). Notably, genetic correlations were approximately twice the magnitude of the phenotypic correlations, and showed similar spatial patterns (**Fig. 5b**). Full results are provided in Supplementary Table 8, with brain maps for each marker in Supplementary Figure 3. To capture the extent of genetic overlap beyond genetic correlations, we used bivariate MiXeR. **Fig. 5c** summarizes analyses with PUFA/MUFA, selected as an exemplar marker given its strong positive genetic correlations with global brain measures, illustrating that the majority of genetic variants influencing this marker overlap with the global brain measures. The concordance rates confirm the presence of mixed effect directions, explaining the lower genetic correlation. Full results are given in Supplementary Table 9.

**Figure 5:**
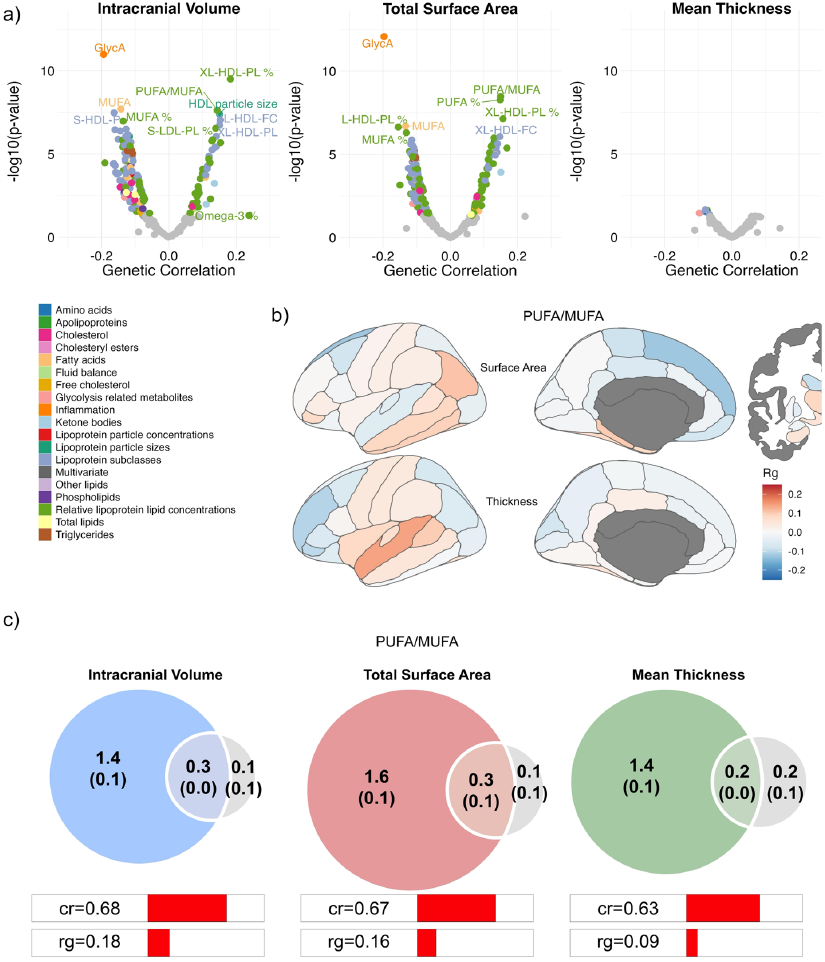
Genetic overlap between individual metabolic markers and brain morphology. **a)** Summary of genetic correlations (r_g_) between each of the 249 metabolites and global brain measures. The x-axis shows the r_g_ estimate and the y-axis shows significance as -log10(p-value). **b)** Brain maps showing the spatial pattern of genetic correlation for an exemplar marker (PUFA/MUFA). **c)** Venn diagrams from bivariate MiXeR illustrating the estimated number of shared and unique causal variants (in thousands) between PUFA/MUFA and global brain measures. Concordance rate (cr) indicates the proportion of shared variants with the same effect direction.

### Shared genetic loci and pathways, and tissue- and cell-type specificity

We ran conjunctional false discovery rate analyses (cFDR)^42^ on each global brain measure paired to each metabolic marker, to identify specific shared genetic variants. We mapped shared lead variants to protein-coding genes using OpenTargets,^43^ and tested these for enrichment among Gene Ontology (GO) biological processes (Supplementary Tables 10,11). The resulting sets of shared genes showed both overlap and specificity across the three global brain measures (**Fig. 6a**). **Fig. 6b** lists the 5 most significantly enriched GO terms for each measure among the 249 marker pairings; for example, shared genes implicated in SA were most enriched in “Stem Cell Differentiation” and the “JNK Cascade”.

**Figure 6:**
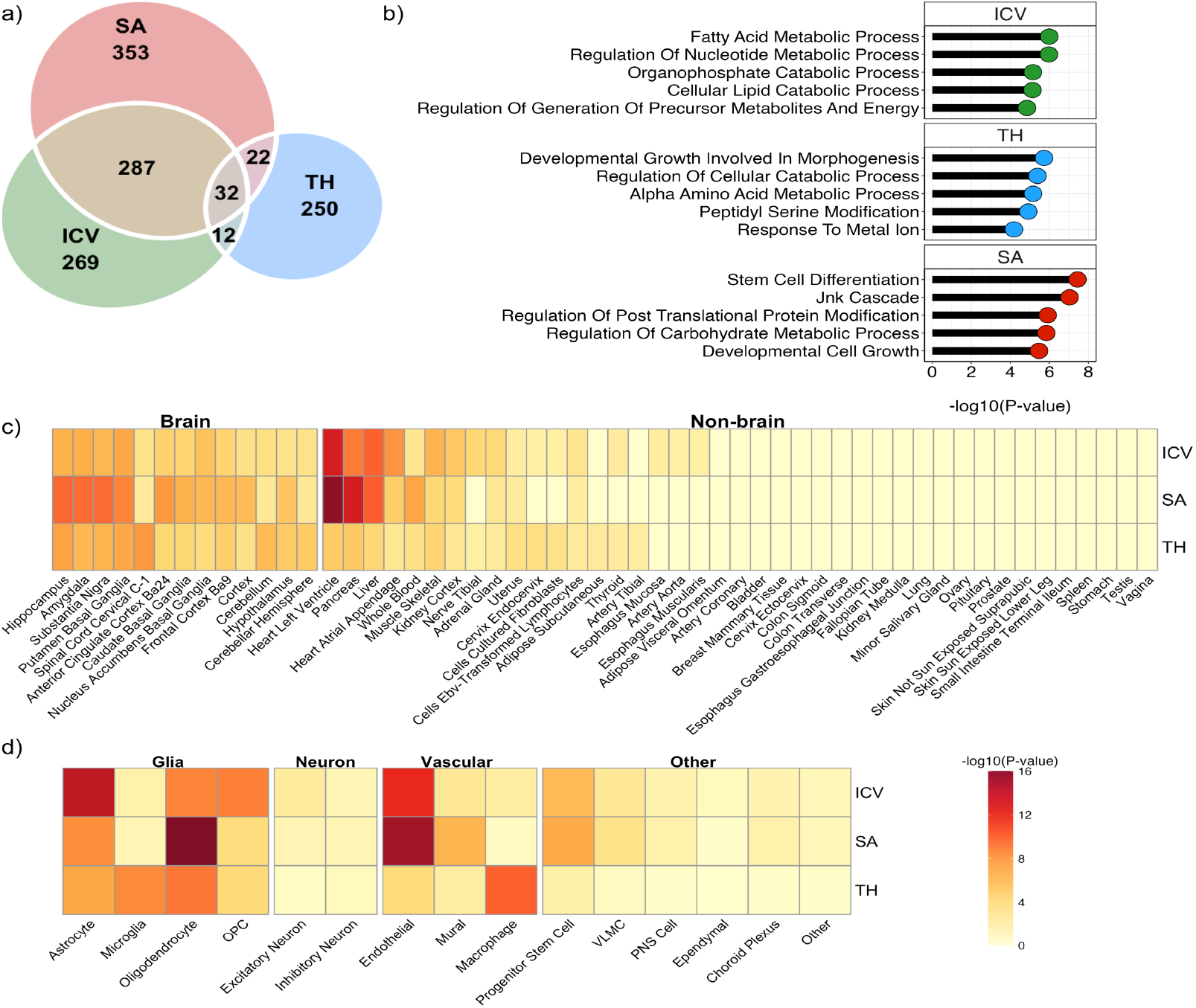
Shared genes, pathways, and tissue enrichment. **a)** Venn diagram showing the number of unique and shared genes identified via conjunctional analysis, aggregated across all metabolites for each global brain measure. **b)** Top 5 enriched Gene Ontology (GO) biological process terms (y-axis) for the shared gene sets with the observed enrichment -log10(p-values) on the x-axis, per global brain measure (panels). **c) and d)** Heatmaps of tissue- and cell type-specific gene expression enrichment. Color intensity reflects the Bonferroni-corrected enrichment significance (-log10(p-value)). Rows are global brain metrics. For **c)**, columns are 54 human tissues from GTEx v8 data, grouped into ‘brain’ and ‘non-brain’ categories. for **d)**, columns are cell types with data taken from 5 different single cell RNAseq human and murine datasets. ICV, intracranial volume; SA, total surface area; TH, mean cortical thickness.

To investigate the tissue-specific expression of these genes shared between systemic metabolism and global brain morphology, we performed enrichment analyses using GTEx v8 data. Genes linked with all three global brain measures exhibited widespread enrichment across numerous brain regions, including the hippocampus, amygdala, various basal ganglia structures, and cortical areas. Notably, exceptionally strong enrichments were also observed in key peripheral tissues: the heart, pancreas, and liver (**Fig. 6c;** Supplementary Table 12). To identify the specific cell types underlying these genetic associations, we performed gene-set enrichment analysis using five independent scRNA-seq datasets (**Figure 6d**; Supplementary Table 13). Aggregating across all metabolites, thereby testing the link with systemic metabolism, genes shared with ICV and SA showed the most powerful enrichment in oligodendrocytes (SA: p = 2.61×10^-24^) and astrocytes (ICV: p=1.22×10^-15^). In contrast, genes shared with TH showed a distinct pattern, with the strongest enrichments in immune cells like macrophages (p=3.49×10^-11^) and microglia (p = 1.89×10^-09^). For GlycA specifically, shared genes were most enriched in astrocytes for ICV (p=5.34×10^-09^) and in microglia for SA (p=2.76×10^-05^), supporting a link between inflammation and glial function. For PUFA/MUFA, shared genes were strongly enriched in myelinating cells, namely oligodendrocytes (SA: p=6.11×10^-05^) and their precursors (ICV: p=1.20×10^-05^).

## Discussion

In this study of the human brain-body metabolic axis, we integrated deep plasma metabolomics with multimodal neuroimaging and genomics to reveal extensive links between systemic metabolism and *in vivo* brain morphology. Our findings provide evidence consistent with widespread, bidirectional causal relationships, identifying specific metabolic markers and pleiotropic gene pathways that act across both brain and key peripheral organs. These results suggest that systemic metabolic markers could help define brain health trajectories, and may serve as targets for interventions to reduce the negative interplay between cardiometabolic and brain dysfunction with ageing.

Systemic metabolism was most strongly linked to global measures of brain morphology, particularly ICV and SA. These measures are determined by developmental mechanisms distinct from those influencing cortical thickness.^38,44^ This was mirrored in the genetic data, where the strongest genetic correlations were between markers like GlycA and PUFA/MUFA and these same two global brain measures. While these brain metrics are established by development, the associations observed in this older cohort likely reflect a lifelong effect of their underlying biological pathways. For example, the critical role of lipid metabolism is well-established in neurodevelopment, where disruptions can lead to conditions like micro- or macrocephaly.^45^ Since the BBB limits peripheral lipoprotein entry, the developing brain depends on local cholesterol synthesis and astrocyte-secreted lipoproteins for axonal growth and membrane biogenesis.^16,46^ Our findings suggest that this fundamental biological axis remains important for maintenance of brain structure throughout adulthood.

Our genetic and cell-type analyses provide a biological context for these observations. Pathway analysis of the shared genes for ICV and surface area converged on fundamental processes like “stem cell differentiation” and the JNK cascade, a pathway with an established dual role in mediating metabolic stress and regulating neural stem cell fate.^47^ Furthermore, these shared genes were most powerfully enriched in oligodendrocytes and astrocytes, which aligns with the finding that these same lipid-dependent processes influence neural progenitor cell fate, a mechanism critical for the evolutionary expansion of the neocortex.^48^

Tissue-specific gene expression analyses further revealed that these shared genetic factors act pleiotropically. Genes linked to ICV and surface area showed enrichment not only in diverse brain regions but also in key peripheral metabolic organs including the liver, pancreas, and heart. This provides genetic evidence that the observed brain-body associations are driven by pleiotropy, likely underpinned by fundamental signaling pathways known to coordinate the development and function of these diverse organ systems.^49^

Our multivariate analyses further clarified the structure of the link between systemic inflammation, lipid metabolism, and brain morphology. The canonical correlation analysis identified a primary component linking larger global and cortical measures to lower systemic inflammation, as indexed by the inflammatory marker GlycA, and higher HDL concentrations. This pattern was consistent across both phenotypic and genetic correlations. Higher GlycA, a biomarker previously found to index aging,^50^ was associated with smaller brain volumes, consistent with the detrimental effects of chronic inflammation on brain integrity.^51^ The opposing association with higher HDL aligns with its established neuroprotective roles.^52^ Since large molecules like GlycA and HDL do not readily cross the blood-brain barrier, their effects on the brain are likely indirect. Our findings support a mechanism whereby systemic inflammation compromises the neurovascular unit or promotes pro-inflammatory glial activation.^53^ Bolstering this interpretation, our cell-type-specific analysis revealed that the genes linking GlycA to brain size are significantly enriched in astrocytes and microglia, key mediators of neuroinflammatory processes.

Beyond global brain size, systemic metabolism was associated with regional brain structure, with the strongest effects observed for cortical thickness. We identified the superior temporal and frontal lobes as primary sites for these associations, regions known for their high metabolic rates and vulnerability to age-related atrophy and glucose hypometabolism.^54,55^ Our Mendelian randomization analyses support this interpretation, providing causal evidence that metabolites influence frontal and temporal cortical thickness. The strong association with the inflammatory marker GlycA, for example, points towards mechanisms such as microglial activation or neurovascular dysfunction contributing to these processes.^56^ The overarching complexity of this axis is underscored by our Mendelian randomization results. We found evidence for consistent negative feedback loops between apolipoprotein A1 (a key HDL component) and multiple frontal cortical regions, pointing to a tightly regulated relationship.

Finally, our analyses revealed a complex association pattern characteristic of brain aging. The second major component of shared variance linked a pattern of smaller ventricles and larger subcortical volumes, indicative of brain structural integrity, to higher concentrations of LDL cholesterol. This finding supports the hypothesis that, in older individuals, ongoing neurodegenerative processes may disrupt the brain’s regulation of systemic cholesterol.^23,57^ Our MR results reinforce this direction of causality, suggesting that variation in subcortical volumes can causally influence plasma metabolite concentrations. This suggests that in an aging population, plasma LDL levels may not only act as a causal risk factor but may also serve as a biomarker of brain structural integrity. Our discovery of an antagonistic loop between small LDL cholesterol and cuneus thickness, where higher LDL was causal for a thicker cuneus which in turn was causal for lower LDL, further highlights this complex regulatory feedback.

Our observation that genetic correlations often exceeded phenotypic ones suggests the full extent of shared biology may be underestimated. The genetic correlation metric can conceal substantial polygenic overlap if the shared variants have a low concordance of effect directions.^58^

Key strengths of the study include our large sample size, comprehensive data, and integrative design, providing a valuable genotype-phenotype atlas. However, some limitations require consideration. The 8-year time lag between metabolic and MRI assessments precludes analysis of short-term dynamics, though our findings likely reflect long-term metabolic influences and our genetic analyses are robust to this separation. Plasma metabolite concentrations are not highly correlated with CNS levels,^59^ meaning many observed effects are likely indirect. We did not test for interactions with age, which may obscure interesting dynamic aspects, as suggested by animal models.^60^ Nor did we covary for the influence of factors such as lifestyle, diet, or clinical comorbidities and medications, which represent potential confounders and mediators that require further investigation. Further, while we employed a robust multi-method MR approach, confounding via unmeasured horizontal pleiotropy cannot be fully excluded, particularly for the indicated bidirectional findings; these specific identified pathways require further validation. Finally, while robust within the large-scale UK Biobank sample, these specific findings await necessary replication in independent cohorts with more diverse ancestries.

In conclusion, by integrating plasma metabolomics with large-scale neuroimaging and multi-modal genomics, we delineate specific metabolic pathways, particularly those governing lipid homeostasis and systemic inflammation, that causally influence human brain morphology. Our findings establish a complex bidirectional axis, where systemic metabolic state impacts brain structure partly through indirect effects on vascular and inflammatory pathways; brain regions, in turn, affect peripheral metabolism. The extensive genetic overlap, acting through pleiotropic effects in both the brain and key metabolic organs like the liver and pancreas, underscores the integrated nature of this relationship. This work moves beyond correlation to characterize the shared molecular landscape of the brain-body metabolic axis, offering a rich atlas of associations that can help prioritize targets for future mechanistic and therapeutic research.

## Supporting information

Supplements Overview

## Methods

A schematic overview of the study design and analytical pipeline is provided in Supplementary Fig. 1. Study sample

We used data from the UK Biobank (UKB) under accession number 27412. The UKB study has received ethics approval from the UK National Health Service Research Ethics Service (ref 11/NW/0382), and all participants provided informed consent. Full study protocols are described elsewhere.^61^ Our final sample consisted of 24,940 unrelated (KING cut-off 0.0884)^62^ participants of White European ancestry with complete metabolomic, neuroimaging, and covariate data (mean age at scan=64.0 years (SD=7.5); mean age at metabolite measurement=55.2 (SD=7.9); 52.5% female).

### Data preprocessing

#### Neuroimaging Data

We processed T1-weighted brain MRI data using the Freesurfer v5.3 default recon-all pipeline. For our primary analyses, we extracted 88 regional and 3 global brain morphology measures. From the cortical stream, we used 34 regional cortical thickness and surface area measures per hemisphere from the Desikan-Killiany atlas,^40^ along with global mean cortical thickness and total surface area. From the subcortical stream, we selected 20 regional volumes and global intracranial volume (ICV).^41^ To increase signal-to-noise and reduce the multiple testing burden, we summed left and right hemisphere measures for all bilateral regions.

We excluded participants with low-quality scans, defined as an age- and sex-corrected Euler number more than 3 standard deviations below the scanner-specific mean. All brain measures were then pre-residualized for sex, age at blood sample collection (baseline), age at scan, scanner site, and Euler number. For regional measures, we also regressed out the corresponding global measure: mean cortical thickness for regional thickness, total surface area for regional area, and ICV for subcortical volumes. We did not include lifestyle or clinical variables (e.g., smoking, BMI, diabetes) as covariates, as these may represent mediators of metabolite–brain associations rather than confounders. However, their influence cannot be fully excluded.

#### Metabolomics Data

We used the 249 plasma metabolic markers from the Nightingale Health NMR panel, which includes 228 lipid-related traits (lipoproteins, cholesterol, fatty acids) and 21 non-lipid traits (e.g., amino acids, glycolysis metabolites, inflammatory markers).^4^ We applied further preprocessing to the UKB-released data using the ‘ukbnmr’ R package to remove technical variation.^63^

Finally, we applied a rank-based inverse normal transformation to all residualized brain measures and all metabolic markers to ensure a normal distribution for downstream analyses.^64^

#### Statistical Analyses

All statistical analyses were conducted in R v4.4. We used the Benjamini-Hochberg procedure to control the false discovery rate (FDR) for multiple comparisons across all 249 metabolites and 91 brain measures, with a significance threshold of FDR<0.05.

#### Phenotypic Association Analyses

We performed linear regression of each pre-residualized brain measure on each metabolic marker. To assess the joint effect of all metabolites on each brain measure, we used partial least squares (PLS) regression via the ‘pls’ R package, extracting the maximum R^2^ from the model output.

#### Canonical Correlation Analysis

We applied CCA to identify multivariate modes of maximum correlation between the set of metabolic markers and the set of regional brain measures. We extracted the first two dominant, orthogonal CCA modes and computed loadings for each feature, calculated as the Pearson correlation between the feature and its corresponding canonical variate.

### Genetic Analyses

#### GWAS

To avoid sample overlap, we first conducted GWAS for all brain and metabolic markers on UKB participants of White European ancestry, using PLINK2 with the same pipeline as used previously.^65^ For brain morphology, GWAS was performed in 39,098 participants. For metabolic markers, GWAS was performed in a non-overlapping sample of 197,475 participants. All GWAS were adjusted for age, sex, scanner, and the first 20 genetic principal components.

#### Mendelian Randomization

We performed bidirectional two-sample MR using the ‘TwoSampleMR’ R package. We selected genome-wide significant variants (P<5×10^-8^) as instrumental variables. We report Inverse Variance Weighted (IVW) MR results and used weighted median^66^ and MR-Egger^67^ methods to check for robustness of findings.

#### Bivariate LDSC

We applied cross-trait LDSC^68^ to estimate genetic correlations between the brain measures and metabolic markers. For this, we formatted the GWAS summary statistics in accordance with recommendations, including ‘munging’ and removal of all variants in the extended major histocompatibility complex (MHC) region (chr6:25–35 Mb).

#### MiXeR

We applied a Gaussian mixture model, MiXeR,^69,70^ to quantify global genetic architectures and estimate the number of shared causal genetic variants between the three global brain measures and metabolic markers. Bivariate MiXeR models additive genetic effects, based on the GWAS summary statistics, as a mixture of four components, representing null SNPs in both traits (π_0_); SNPs with a specific effect on the first and on the second trait (π_1_ and π_2_, respectively); and SNPs with non-zero effect on both traits (π_12_;, please see ^69,70^ for the mathematical framework). We checked whether the models generated positive AIC values compared to a baseline infinitesimal model, indicating good model fit. All 249 metabolic markers and 88 out of the 91 brain measures passed the univariate AIC check, all bivariate models passed the quality checks; see Supplementary Tables 7 and 9.

#### Conjunctional FDR

We conducted conjunctional false discovery rate (cFDR) analyses for each of the brain measures with each of the 249 markers. The cFDR analyses consisted of conditioning the GWAS summary statistics for each of the pair of traits onto each other in both directions through the pleioFDR tool using default settings, see https://github.com/precimed/pleiofdr, whereby the highest of the resulting pair of FDR values per genetic variant was taken as the strength of evidence for its shared effects.^42^ We set an FDR<0.05 threshold, in accordance with recommendations.

#### Gene mapping

We used the Variant-to-Gene (V2G) pipeline from Open Targets Genetics, to map lead variants from the cFDR analyses to genes based on the strongest evidence from quantitative trait loci (QTL) experiments, chromatin interaction experiments, in silico functional prediction, and proximity of each variant to the canonical transcription start site of genes.^43^

#### Gene set enrichment

We carried out gene set enrichment analyses on the cFDR mapped genes, investigating terms that are part of the Gene Ontology biological processes subset (n=7522), as listed in the Molecular Signatures Database (MsigdB; c5.bp.v7.1). We ran Fisher’s exact tests on the mapped genes from each individual cFDR analysis, leveraging the enricher R package. We restricted the output to those terms that had more than 5 overlapping genes and that were smaller than 500 genes in total, with a q-value <.05. We further removed terms from the list of significant enrichments if more than 80% of their overlapping genes were among more significant terms. Running the enrichment analyses on individual cFDR sets of genes, with the above settings, was chosen to ensure that we identified more biologically specific terms than when running across aggregated gene lists and included larger pathways.

#### Expression specificity analyses

To investigate tissue specificity, we tested for enrichment of the cFDR mapped genes among gene sets representing 54 tissues from the GTEx v8 database,^71^ using enrichment analyses through Fisher’s exact tests. To identify the specific cell types contributing to the shared genetic architecture, we performed gene-set enrichment analysis using five independent scRNA-seq datasets from adult human and murine brain tissue.^72–76^ For each global brain measure (ICV, SA, and TH), we tested the corresponding sets of shared genes from the cFDR analyses for enrichment in cell-type-specific gene lists derived from these datasets. To generate a single summary statistic, results from the five datasets were aggregated using Fisher’s method for combining p-values for each major cell type. Significance was determined using a false discovery rate (FDR) threshold of q <.05.

## Acknowledgements

This work was partly performed on the TSD (Tjeneste for Sensitive Data) facilities, owned by the University of Oslo, operated and developed by the TSD service group at the University of Oslo, IT-Department (USIT). Computations were also performed on resources provided by UNINETT Sigma2 - the National Infrastructure for High-Performance Computing and Data Storage in Norway (NS9666S).

## Materials & Correspondence

The data incorporated in this work were gathered from public resources. The code is available via https://github.com/precimed (GPLv3 license). Correspondence and requests for materials should be addressed to <u>d.v.d.meer[at]medisin.uio.no</u>

## Author contributions

D.v.d.M. and O.A.A. conceived the study; D.v.d.M. and J.K. pre-processed the data. D.v.d.M., J.K. and J.R. performed all analyses, with conceptual input from J.F., A.S., S.S., A.M.D., and O.A.A.; All authors contributed to interpretation of results; D.v.d.M. drafted the manuscript and all authors contributed to and approved the final manuscript.

## References

1. Camandola, S. & Mattson, M. P. Brain metabolism in health, aging, and neurodegeneration. EMBO J. 36, 1474–1492 (2017).

2. Roh, E., Song, D. K. & Kim, M.-S. Emerging role of the brain in the homeostatic regulation of energy and glucose metabolism. Exp. Mol. Med. 48, e216–e216 (2016).

3. He, Y., Xie, K., Yang, K., Wang, N. & Zhang, L. Unraveling the Interplay Between Metabolism and Neurodevelopment in Health and Disease. CNS Neurosci. Ther. 31, e70427 (2025).

4. Julkunen, H. et al. Atlas of plasma NMR biomarkers for health and disease in 118,461 individuals from the UK Biobank. Nat. Commun. 14, 604 (2023).

5. van der Meer, D. et al. Divergent patterns of genetic overlap between severe mental disorders and metabolic markers. medRxiv 2024–11 (2024).

6. He, S., Xu, Z. & Han, X. Lipidome disruption in Alzheimer’s disease brain: detection, pathological mechanisms, and therapeutic implications. Mol. Neurodegener. 20, 11 (2025).

7. Vancampfort, D. et al. Risk of metabolic syndrome and its components in people with schizophrenia and related psychotic disorders, bipolar disorder and major depressive disorder: a systematic review and meta-analysis. World Psychiatry 14, 339–347 (2015).

8. Santiago, J. A. & Potashkin, J. A. The impact of disease comorbidities in Alzheimer’s disease. Front. Aging Neurosci. 13, 631770 (2021).

9. Banks, W. A. The blood-brain barrier: connecting the gut and the brain. Regul. Pept. 149, 11–14 (2008).

10. Saklayen, M. G. The global epidemic of the metabolic syndrome. Curr. Hypertens. Rep. 20, 1–8 (2018).

11. Buergel, T. et al. Metabolomic profiles predict individual multidisease outcomes. Nat. Med. 28, 2309–2320 (2022).

12. Lippert, R. N. & Brüning, J. C. Maternal Metabolic Programming of the Developing Central Nervous System: Unified Pathways to Metabolic and Psychiatric Disorders. Metab. Links Body Brain Behav. 91, 898–906 (2022).

13. Bélanger, M., Allaman, I. & Magistretti, P. J. Brain Energy Metabolism: Focus on Astrocyte-Neuron Metabolic Cooperation. Cell Metab. 14, 724–738 (2011).

14. Falkowska, A. et al. Energy metabolism of the brain, including the cooperation between astrocytes and neurons, especially in the context of glycogen metabolism. Int. J. Mol. Sci. 16, 25959–25981 (2015).

15. Bruce, K. D., Zsombok, A. & Eckel, R. H. Lipid processing in the brain: a key regulator of systemic metabolism. Front. Endocrinol. 8, 60 (2017).

16. Yoon, J. H. et al. Brain lipidomics: From functional landscape to clinical significance. Sci. Adv. 8, eadc9317 (2022).

17. Müller, C. P. et al. Brain membrane lipids in major depression and anxiety disorders. Biochim. Biophys. Acta BBA-Mol. Cell Biol. Lipids 1851, 1052–1065 (2015).

18. Bazinet, R. P. & Layé, S. Polyunsaturated fatty acids and their metabolites in brain function and disease. Nat. Rev. Neurosci. 15, 771–785 (2014).

19. Teleanu, R. I. et al. Neurotransmitters—key factors in neurological and neurodegenerative disorders of the central nervous system. Int. J. Mol. Sci. 23, 5954 (2022).

20. Grimm, A. & Eckert, A. Brain aging and neurodegeneration: from a mitochondrial point of view. J. Neurochem. 143, 418–431 (2017).

21. Kennedy, K. G. et al. Elevated lipids are associated with reduced regional brain structure in youth with bipolar disorder. Acta Psychiatr. Scand. 143, 513–525 (2021).

22. Ward, M. A. et al. Low HDL cholesterol is associated with lower gray matter volume in cognitively healthy adults. Front. Aging Neurosci. 2, 29 (2010).

23. Moazzami, K. et al. Association of mid-life serum lipid levels with late-life brain volumes: The atherosclerosis risk in communities neurocognitive study (ARICNCS). Neuroimage 223, 117324 (2020).

24. Gordon, S. et al. Metabolites and MRI-Derived Markers of AD/ADRD Risk in a Puerto Rican Cohort. Res. Sq. (2024).

25. Kindler, J. et al. Dysregulation of kynurenine metabolism is related to proinflammatory cytokines, attention, and prefrontal cortex volume in schizophrenia. Mol. Psychiatry 25, 2860–2872 (2020).

26. Gurholt, T. P. et al. Population-based body–brain mapping links brain morphology with anthropometrics and body composition. Transl. Psychiatry 11, 1–12 (2021).

27. Beck, D. et al. Cardiometabolic risk factors associated with brain age and accelerated brain ageing. Hum. Brain Mapp. 43, 700–720 (2022).

28. Henkel, N. D. et al. Schizophrenia: A disorder of broken brain bioenergetics. Mol. Psychiatry 27, 2393–2404 (2022).

29. Prabakaran, S. et al. Mitochondrial dysfunction in schizophrenia: evidence for compromised brain metabolism and oxidative stress. Mol. Psychiatry 9, 684–697 (2004).

30. Du, F. et al. In Vivo Evidence for Cerebral Bioenergetic Abnormalities in Schizophrenia Measured Using 31P Magnetization Transfer Spectroscopy. JAMA Psychiatry 71, 19–27 (2014).

31. Zeng, Y., Guo, R., Cao, S. & Yang, H. Causal associations between blood lipids and brain structures: a Mendelian randomization study. Cereb. Cortex 33, 10901–10908 (2023).

32. Lin, B. D. et al. Dissecting causal relationships between cortical morphology and neuropsychiatric disorders: a bidirectional Mendelian randomization study. medRxiv 2024–09 (2024).

33. Rødevand, L. et al. Characterizing the shared genetic underpinnings of schizophrenia and cardiovascular disease risk factors. Am. J. Psychiatry 180, 815–826 (2023).

34. Amare, A. T., Schubert, K. O., Klingler-Hoffmann, M., Cohen-Woods, S. & Baune, B. T. The genetic overlap between mood disorders and cardiometabolic diseases: a systematic review of genome wide and candidate gene studies. Transl. Psychiatry 7, e1007–e1007 (2017).

35. van der Meer, D. et al. Pleiotropic and sex-specific genetic architecture of circulating metabolic markers. medRxiv 2024.07. 30.24311254 (2024).

36. Karjalainen, M. K. et al. Genome-wide characterization of circulating metabolic biomarkers. Nature (2024) doi:10.1038/s41586-024-07148-y.

37. van der Meer, D. et al. Understanding the genetic determinants of the brain with MOSTest. Nat. Commun. 11, 3512 (2020).

38. Grasby, K. L. et al. The genetic architecture of the human cerebral cortex. bioRxiv 399402–399402 (2018) doi:10.1101/399402.

39. Hampel, H. et al. The foundation and architecture of precision medicine in neurology and psychiatry. Trends Neurosci. 46, 176–198 (2023).

40. Desikan, R. S. et al. An automated labeling system for subdividing the human cerebral cortex on MRI scans into gyral based regions of interest. NeuroImage 31, 968–980 (2006).

41. Fischl, B. et al. Whole brain segmentation: automated labeling of neuroanatomical structures in the human brain. Neuron 33, 341–355 (2002).

42. Smeland, O. B. et al. Discovery of shared genomic loci using the conditional false discovery rate approach. Hum. Genet. 139, 85–94 (2020).

43. Mountjoy, E. et al. An open approach to systematically prioritize causal variants and genes at all published human GWAS trait-associated loci. Nat. Genet. 53, 1527–1533 (2021).

44. Winkler, A. M. et al. Cortical thickness or grey matter volume? The importance of selecting the phenotype for imaging genetics studies. NeuroImage 53, 1135–1146 (2010).

45. Xing, L., Huttner, W. B. & Namba, T. Role of cell metabolism in the pathophysiology of brain size-associated neurodevelopmental disorders. Neurobiol. Dis. 199, 106607 (2024).

46. Hussain, G. et al. Role of cholesterol and sphingolipids in brain development and neurological diseases. Lipids Health Dis. 18, 1–12 (2019).

47. de los Reyes Corrales, T., Losada-Pérez, M. & Casas-Tintó, S. JNK pathway in CNS pathologies. Int. J. Mol. Sci. 22, 3883 (2021).

48. Xing, L. et al. Functional synergy of a human-specific and an ape-specific metabolic regulator in human neocortex development. Nat. Commun. 15, 3468 (2024).

49. Foulquier, S. et al. WNT signaling in cardiac and vascular disease. Pharmacol. Rev. 70, 68–141 (2018).

50. Zhang, S. et al. A metabolomic profile of biological aging in 250,341 individuals from the UK Biobank. Nat. Commun. 15, 8081 (2024).

51. Williams, J. A. et al. Inflammation and brain structure in schizophrenia and other neuropsychiatric disorders: a Mendelian randomization study. JAMA Psychiatry 79, 498–507 (2022).

52. Turri, M., Marchi, C., Adorni, M. P., Calabresi, L. & Zimetti, F. Emerging role of HDL in brain cholesterol metabolism and neurodegenerative disorders. Biochim. Biophys. Acta BBA-Mol. Cell Biol. Lipids 1867, 159123 (2022).

53. Jiménez, A. et al. Nanotechnology to Overcome Blood–Brain Barrier Permeability and Damage in Neurodegenerative Diseases. Pharmaceutics 17, 281 (2025).

54. Deery, H. A., Di Paolo, R., Moran, C., Egan, G. F. & Jamadar, S. D. Lower brain glucose metabolism in normal ageing is predominantly frontal and temporal: A systematic review and pooled effect size and activation likelihood estimates meta-analyses. Hum. Brain Mapp. 44, 1251–1277 (2023).

55. Choi, M. et al. Comparison of neurodegenerative types using different brain MRI analysis metrics in older adults with normal cognition, mild cognitive impairment, and Alzheimer’s dementia. PloS One 14, e0220739 (2019).

56. Sweeney, M. D., Sagare, A. P. & Zlokovic, B. V. Blood–brain barrier breakdown in Alzheimer disease and other neurodegenerative disorders. Nat. Rev. Neurol. 14, 133–150 (2018).

57. Solomon, A. et al. Plasma levels of 24S-hydroxycholesterol reflect brain volumes in patients without objective cognitive impairment but not in those with Alzheimer’s disease. Neurosci. Lett. 462, 89–93 (2009).

58. van der Meer, D. et al. Mapping the genetic landscape of psychiatric disorders with the MiXeR toolset. Biol. Psychiatry (2025).

59. Huo, Z. et al. Brain and blood metabolome for Alzheimer’s dementia: findings from a targeted metabolomics analysis. Neurobiol. Aging 86, 123–133 (2020).

60. Ding, J. et al. A metabolome atlas of the aging mouse brain. Nat. Commun. 12, 6021 (2021).

61. Sudlow, C. et al. UK biobank: an open access resource for identifying the causes of a wide range of complex diseases of middle and old age. PLoS Med. 12, e1001779–e1001779 (2015).

62. Manichaikul, A. et al. Robust relationship inference in genome-wide association studies. Bioinformatics 26, 2867–2873 (2010).

63. Ritchie, S. C. et al. Quality control and removal of technical variation of NMR metabolic biomarker data in ∼120,000 UK Biobank participants. Sci. Data 10, 64–64 (2023).

64. Beasley, T. M., Erickson, S. & Allison, D. B. Rank-based inverse normal transformations are increasingly used, but are they merited? Behav. Genet. 39, 580–595 (2009).

65. van der Meer, D. et al. Pleiotropic and sex-specific genetic mechanisms of circulating metabolic markers. Nat. Commun. 16, 1–12 (2025).

66. Bowden, J., Davey Smith, G., Haycock, P. C. & Burgess, S. Consistent estimation in Mendelian randomization with some invalid instruments using a weighted median estimator. Genet. Epidemiol. 40, 304–314 (2016).

67. Burgess, S. & Thompson, S. G. Interpreting findings from Mendelian randomization using the MR-Egger method. Eur. J. Epidemiol. 32, 377–389 (2017).

68. Bulik-Sullivan, B. et al. An atlas of genetic correlations across human diseases and traits. Nat. Genet. 47, 1236 (2015).

69. Holland, D. et al. Beyond SNP Heritability: Polygenicity and Discoverability of Phenotypes Estimated with a Univariate Gaussian Mixture Model. bioRxiv 498550 (2019) doi:10.1101/498550.

70. Frei, O. et al. Bivariate causal mixture model quantifies polygenic overlap between complex traits beyond genetic correlation. Nat. Commun. 10, 2417 (2019).

71. GTEx Consortium. The GTEx Consortium atlas of genetic regulatory effects across human tissues. Science 369, 1318–1330 (2020).

72. Habib, N. et al. Div-Seq: Single-nucleus RNA-Seq reveals dynamics of rare adult newborn neurons. Science 353, 925–928 (2016).

73. Lake, B. B. et al. Integrative single-cell analysis of transcriptional and epigenetic states in the human adult brain. Nat. Biotechnol. 36, 70–80 (2018).

74. Saunders, A. et al. Molecular diversity and specializations among the cells of the adult mouse brain. Cell 174, 1015-1030. e16 (2018).

75. Skene, N. G. et al. Genetic identification of brain cell types underlying schizophrenia. Nat. Genet. 50, 825–833 (2018).

76. Zeisel, A. et al. Molecular architecture of the mouse nervous system. Cell 174, 999–1014 (2018).

