## Supplements Overview for "Shared Molecular Landscape of Human Brain Structure and Systemic Metabolism"

**Atlas of genetic overlap between plasma metabolic markers and brain morphology**

*Supplementary Information*

Here we provide an overview of the manuscript supplementary files, together with explanation of the contents of these files, including the column names.

Overview

Supplementary Table ST1: Information on included brain measures

Supplementary Table ST2: Information on included metabolites

Supplementary Table ST3: Inferential statistics from all univariate linear regressions

Supplementary Table ST4: Inferential statistics from regressions after excluding disorders

Supplementary Table ST5: CCA loadings

Supplementary Table ST6: Mendelian randomization results

Supplementary Table ST7: Global genetic architecture estimates for brain and metabolites

Supplementary Table ST8: Genetic correlations

Supplementary Table ST9: Bivariate MiXeR output

Supplementary Table ST10: List of conjunctional FDR loci and mapped genes

Supplementary Table ST11: Enriched Gene Ontology terms

Supplementary Table ST12: Tissue-specific gene expression results

Supplementary Table ST13: Cell type-specific gene expression results

(All Supplementary Tables are provided as separate Excel files)

Supplementary Figure SF1: Study overview figure

Supplementary Figure SF2: Brain maps for all metabolites – phenotypic association

Supplementary Figure SF3: Scatterplot of global genetic architecture measures

Supplementary Figure SF4: Brain maps for all metabolites – genetic correlation

(SF2 and SF4 are provided as separate zip files, for ease of handling of 249 individual maps)

*Supplementary Table ST1: Information on included brain measures*

This table contains an overview of all brain measures included in the study, generated by applying Freesurfer to the UK Biobank T1 brain MRI data. It contains the following columns:

‘origName’ = original name of the brain measure (as provided by Freesurfer output); ‘prettyName’ = manually formatted brain measure name; ‘metric’ = the type of brain measure (either ‘aseg’ for subcortical, ‘thickness’ for cortical thickness, ‘area’ for surface area, or ‘global’ for one of the three global brain measures); ‘global’ = indicator of which global measure was used to regress out effects of global brain size; ‘lobe’= indicator of which cortical lobe the brain measure is considered to be a part of.

*Supplementary Table ST2: Information on included metabolites*

This table contains an overview of all metabolic markers included in the study, as provided by the UK Biobank, with the following columns:

‘fullName’ = metabolic marker; ‘prettyName’ = formatted marker name; ‘nmrName’= marker name used by the Nightingale QC pipeline; ‘clinicalMarker’ = whether the marker has been clinically validated by Nightingale (‘0’ is no, ‘1’ is yes); ‘group’ = biological subgroup of the marker; ‘groupColor’ = color used for plotting.

*Supplementary Table ST3: Inferential statistics from all univariate linear regressions*

This table provides the full inferential statistics extracted from each univariate regression, whereby each (pre-residualized) brain region was regressed onto each individual marker (i.e. 91 * 249 univariate models). It contains the following columns:

‘marker’ = the metabolic marker as predictor; ‘feature’ = the brain measure as outcome; ‘beta’= regression coefficient; ‘se’= standard error; ‘pL’= p-value; ‘p_adjust’= p-value corrected for multiple comparisons through the Benjamini-Hochberg method; ‘zval’= z-value; ‘r2’= r-squared value of the model.

*Supplementary Table ST4: Inferential statistics from regressions after excluding disorders*

This table provides the full inferential statistics extracted from each univariate regression, after excluding individuals with an ICD10 G2 or G3 diagnosis. Each (pre-residualized) brain region was regressed onto each individual marker (i.e. 91 * 249 univariate models). It contains the following columns:

‘marker’ = the metabolic marker as predictor; ‘feature’ = the brain measure as outcome; ‘beta’= regression coefficient; ‘se’= standard error; ‘pL’= p-value; ‘p_adjust’= p-value corrected for multiple comparisons through the Benjamini-Hochberg method; ‘zval’= z-value; ‘r2’= r-squared value of the model.

*Supplementary Table ST5: CCA loadings*

This table provides the loadings extracted from the canonical correlation analyses, whereby the first column (‘feature’) indicates either the marker or the brain measure. The remaining ten columns indicate the specific component (‘C1’ through ‘C10’), with the cells containing the loadings.

*Supplementary Table ST6: Mendelian randomization results*

These tables (‘a’ and ‘b’) lists the results of the Mendelian randomization analyses, in both directions (metabolites onto brain, and brain onto metabolites, resp.). They contain the following columns:

‘brain’ = brain measure under investigation; ‘marker’ = metabolic marker under investigation ‘beta’= regression coefficient; ‘se’= standard error; ‘pval’= p-value; ‘method’ = MR method.

*Supplementary Table ST7: Global genetic architecture estimates for brain and metabolites*

This table contains the output of the MiXeR analyses, quantifying global genetic architecture of the brain and metabolite measures. It contains the following columns:

‘measure’ = brain or metabolic marker under investigation; ‘h2’ = the estimated SNP-based heritability; ‘h2_sd’= the standard deviation of the heritability estimate across twenty runs; ‘discoverability’ = σ^2^_β_, the variance of effect sizes of non-null genetic variants; ‘disc_sd’= the standard deviation of the discoverability estimate across twenty runs; ‘polygenicity’ = nc@p9, the number of non-null variants to explain 90% of h^2^_snp_; ‘poly_sd’= the standard deviation of the polygenicity estimate across twenty runs.

*Supplementary Table ST8: Genetic correlations*

This table contains the inferential statistics of the genetic correlations between the GWAS summary statistics of the metabolic markers and the brain measures, as calculated through LD score regression. It has the following columns:

‘marker’ = metabolic marker under investigation; ‘rg’ = genetic correlation between the marker and the trait; ‘se’ = the accompanying standard error; ‘zval’= associated z-value; ‘p’= associated p-value.

*Supplementary Table ST9: Bivariate MiXeR output*

This table contains the output of the bivariate MiXeR analyses, between selected metabolites and the three global brain measures. It contains the following columns:

‘marker’ = metabolic marker under investigation; ‘brainMeasure’ = the brain measure; ‘dice’=the Dice coefficient of the overlap, ‘poly1’=the polygenicity estimate of the marker; ‘poly2’=the polygenicity estimate of the brain measure; ‘overlap’ = the estimated number of shared variants between the metabolite and the brain measure; ‘*_sd’; the standard deviation of the estimates across twenty runs; ‘concordance’ = concordance of effect direction of shared variants.

*Supplementary Table ST10: List of conjunctional FDR loci and mapped genes*

This table lists the loci discovered through the conjunctional FDR analyses between the metabolites and the brain measures. It contains the following columns:

‘marker’ = metabolic marker under investigation; ‘brainMeasure’=the brain measure; ‘locusnum’ = index of the loci discovered; ‘SNP’ = identified lead variant;	‘CHR’= chromosome; ‘BP’ = lead variant position on the chromosome in base pairs; ‘conjfdr’=conjunctional FDR value; ‘gene’ = gene mapped to the lead variant through OpenTargets.

*Supplementary Table ST11: Enriched Gene Ontology terms*

This table list significant Gene Ontology biological processes that were found through the enrichment tests on the sets of genes mapped to the conjunctional FDR-discovered lead variants for the global brain measures. The table has the following columns:

‘brainMeasure’=the brain measure; ‘metabo’= metabolic marker under investigation; ’term’ = significant Gene Ontology biological process; ‘GeneRatio’ = number of genes in the GO term found to be overlapping with the cFDR set divided by the total number of genes in the cFDR set; ‘BgRatio’ = number of genes in the GO term divided by the total number of genes, i.e. all protein-coding genes; ‘pvalue’ = p-value of the enrichment test; ‘qvalue’ = q-value of the enrichment test.

*Supplementary Table ST12: Tissue-specific gene expression results*

This table provides the output from tissue-specificity analyses, using Fisher’s test to check for enrichment of genes overlapping between global brain measures and metabolites (mapped genes from the cFDR analyses), among 54 tissue types that are part of the GTEx dataset. It has the following columns:

‘GeneSet’=the tissue type under investigation; ‘N_genes’= number of genes differentially expressed in the tissue; ’ N_overlap’ = number of genes overlapping between the tissue and the mapped gene list; ‘p’ = p-value from the Fisher’s test; ‘adjP’ = p-value adjusted for multiple comparisons; ‘BrainMetric’ = the global brain measure investigated; ‘Proportion’ = proportion of overlapping genes compared to total number of genes expressed in the tissue; ‘TissueGroup’ = category that the tissue type belongs to, brain or non-brain.

*Supplementary Table ST13: Cell type-specific gene expression results*

This table provides the output from cell type-specificity analyses, using Fisher’s test to check for enrichment of genes overlapping between global brain measures and metabolites (mapped genes from the cFDR analyses), among five single-cell RNAseq datasets. It has the following columns:

‘brain’=the brain measure; ‘metabolite’= metabolic marker under investigation; ’dataset’ = scRNAseq dataset tested; ‘cellType’ = cell type under investigation; ‘nOverlapGenes’ = number of genes overlapping between the cell type and the mapped gene list; ‘pval’ = p-value of the enrichment test; ‘OR’ = odds ratio; ‘FDR’ = FDR-value of the enrichment test.

*Supplementary Figure SF1: Study overview*

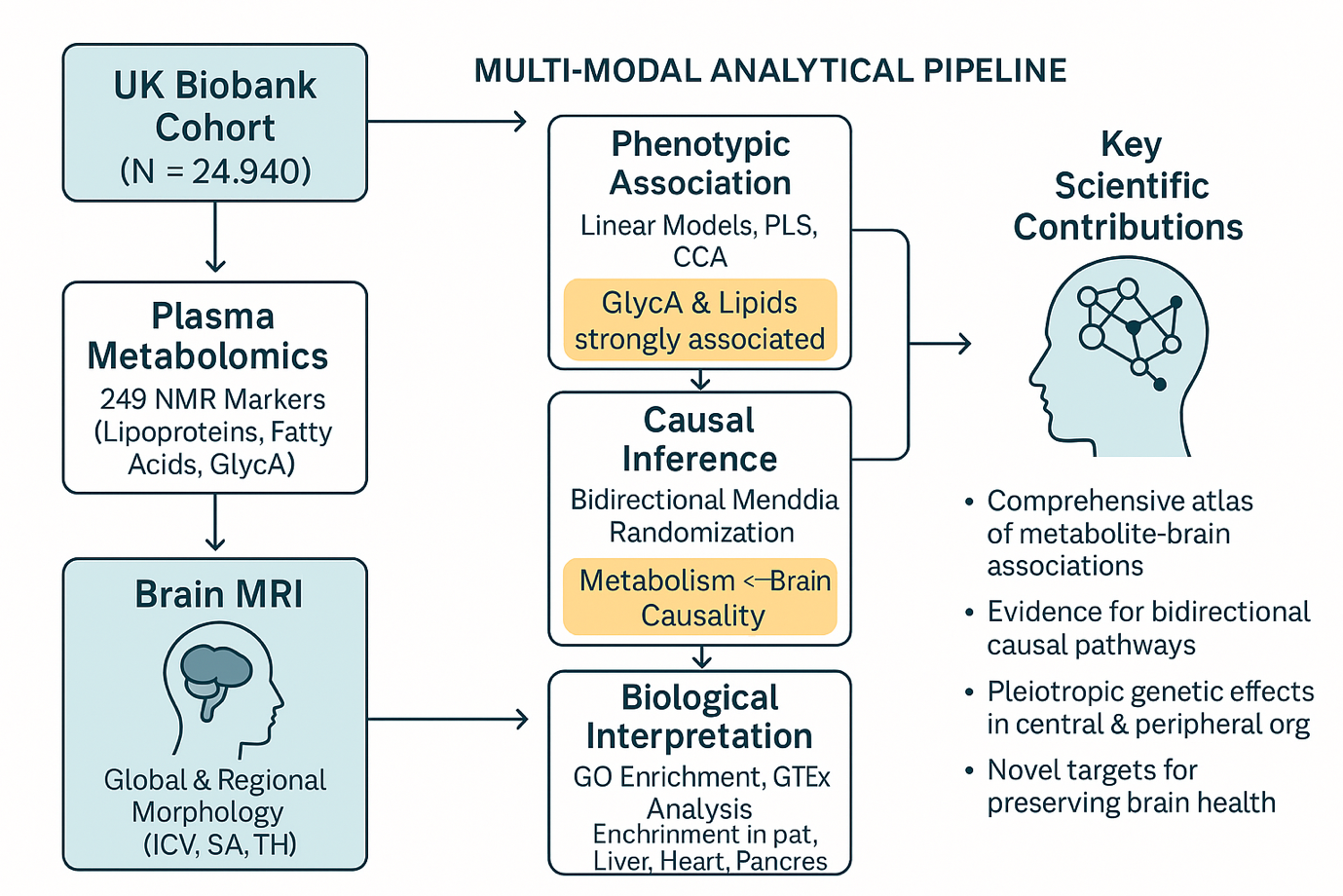

*Supplementary Figure SF2: Brain maps for all metabolites – phenotypic association*

These brain maps, provided in a separate zip file due to the number of maps and their file size, illustrate the strength of association between the markers and brain measures, by depicting the estimated z-scores from linear regression analyses for each brain region. There is a map for each individual marker. The caption is as follows:

*Brain maps depicting the strength of association (z-values, color-coded) from linear regression with the metabolic marker (indicated at the top) as predictor and the different regional brain measures as outcomes, thereby showcasing spatial patterns. The top left brain maps depict the lateral and medial view of the regional surface area results, the bottom left maps depict the regional cortical thickness results, and the brain map in the top right indicates the results for subcortical volumes.*

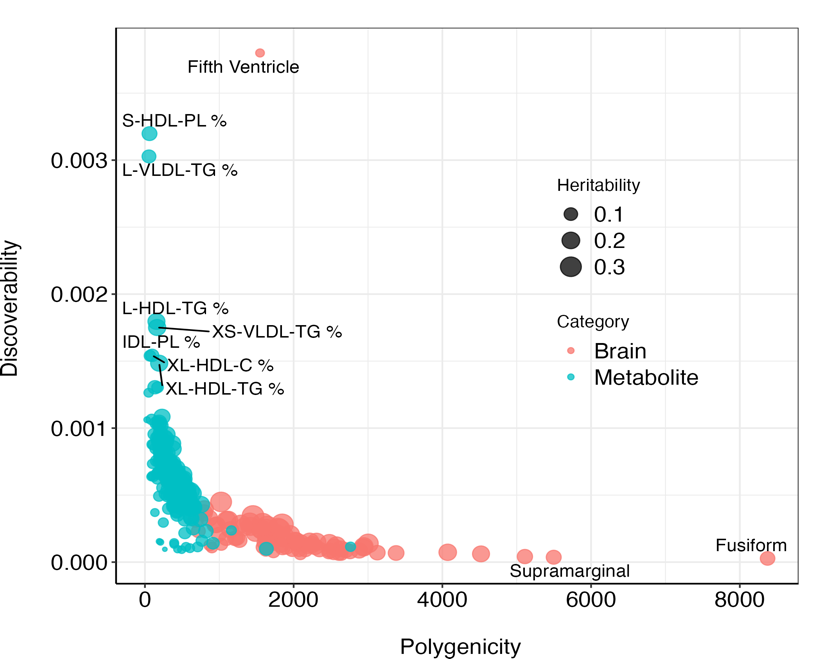

*Supplementary Figure SF3: Scatterplot of global genetic architecture measures*

*Scatterplot summarizing measures of global genetic architecture for metabolites (in green) and brain (in red). The discoverability is on the y-axis, the polygenicity on the x-axis. The size of the dots reflect their heritability. Measures with values at the end of the distributions are labeled.*

*Supplementary Figure SF4: Brain maps for all metabolites – genetic correlation*

These brain maps, provided in a separate zip file due to the number of maps and their file size, depict the estimated genetic correlations from linkage disequilibrium score regression analyses for each brain region. There is a map for each individual marker. The caption is as follows:

*Brain maps depicting the genetic correlation (rg, color-coded) between the metabolic marker (indicated at the top) and the different regional brain measures as outcomes, thereby showcasing spatial patterns. The top left brain maps depict the lateral and medial view of the regional surface area results, the bottom left maps depict the regional cortical thickness results, and the brain map in the top right indicates the results for subcortical volumes.*
